## Supplementary figures and images for "EROS is a selective chaperone regulating the phagocyte NADPH oxidase and purinergic signalling"

### Figure 1- figure supplement 1

**Figure 1 - Figure supplement 1**

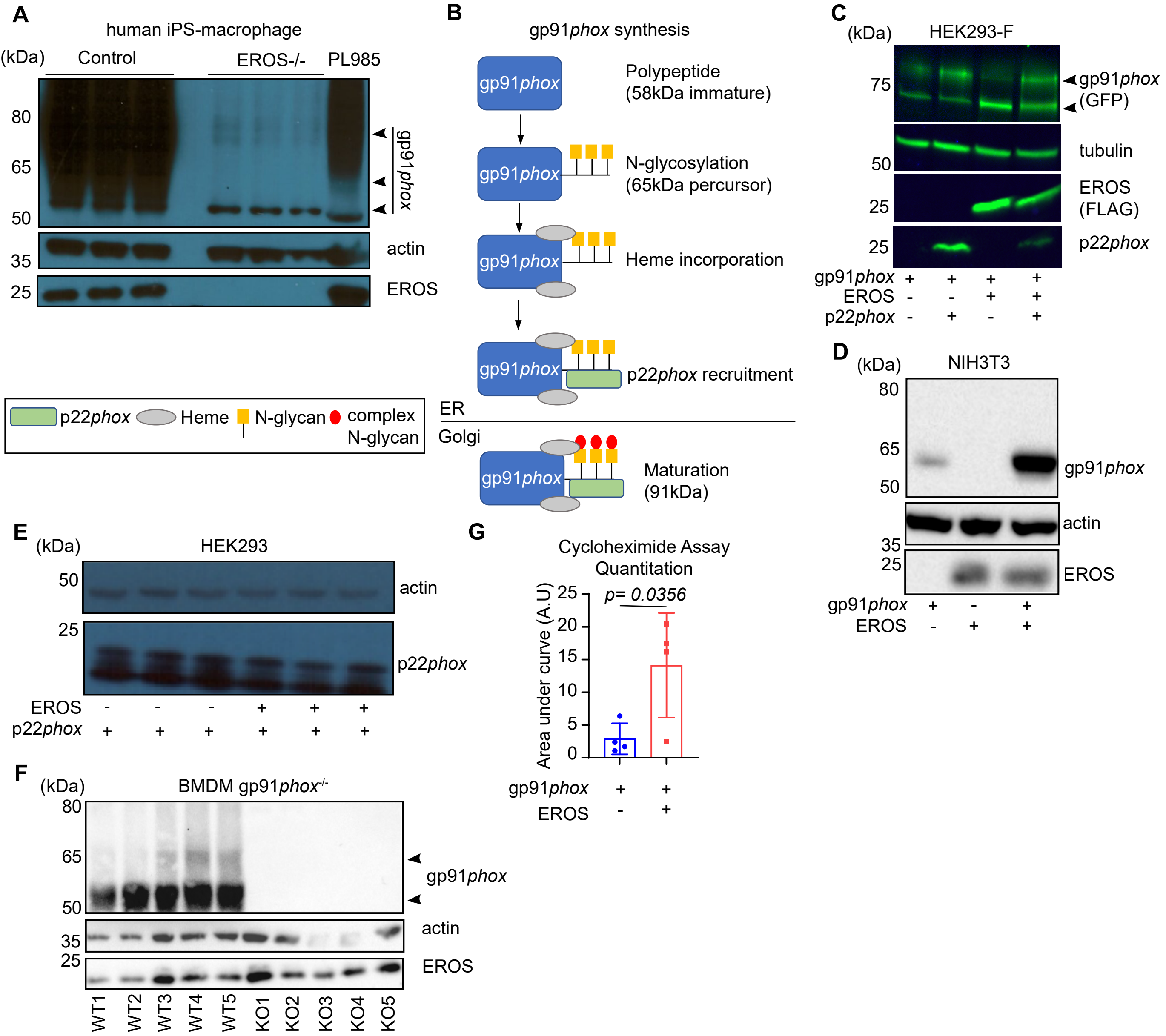

### Figure 2- figure supplement 1

**Figure 2 - Figure supplement 1**

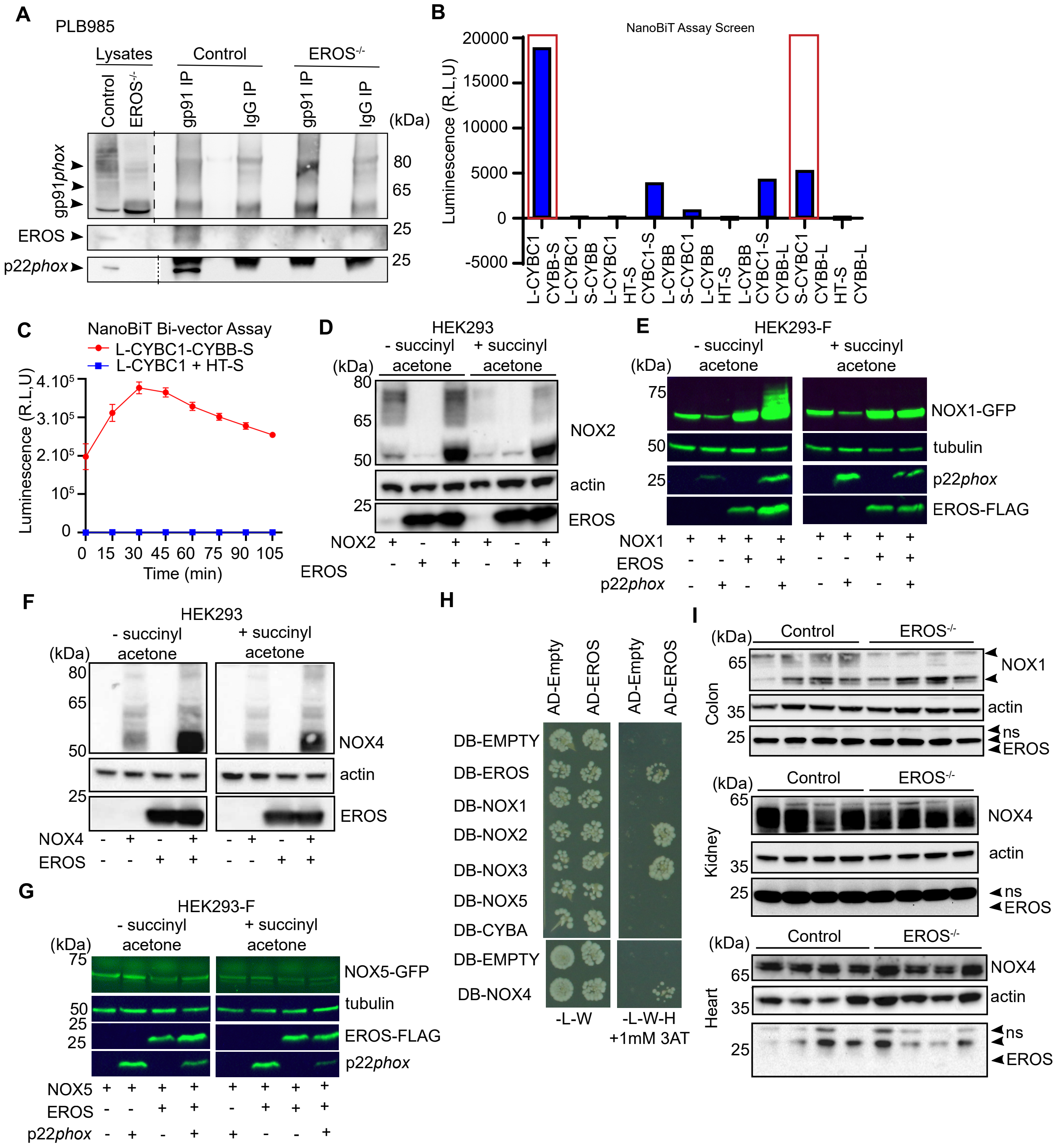

### Figure 2- figure supplement 2

**Figure 2 - Figure supplement 2**

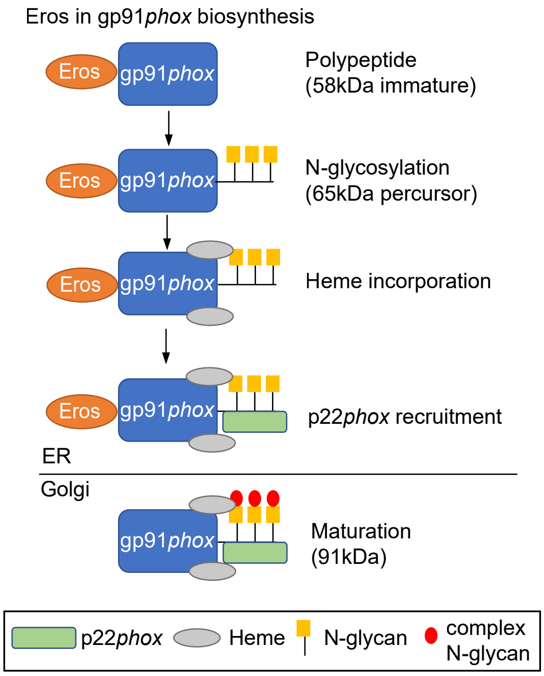

### Figure 3- figure supplement 1

#### Figure 3 - Figure supplement 1

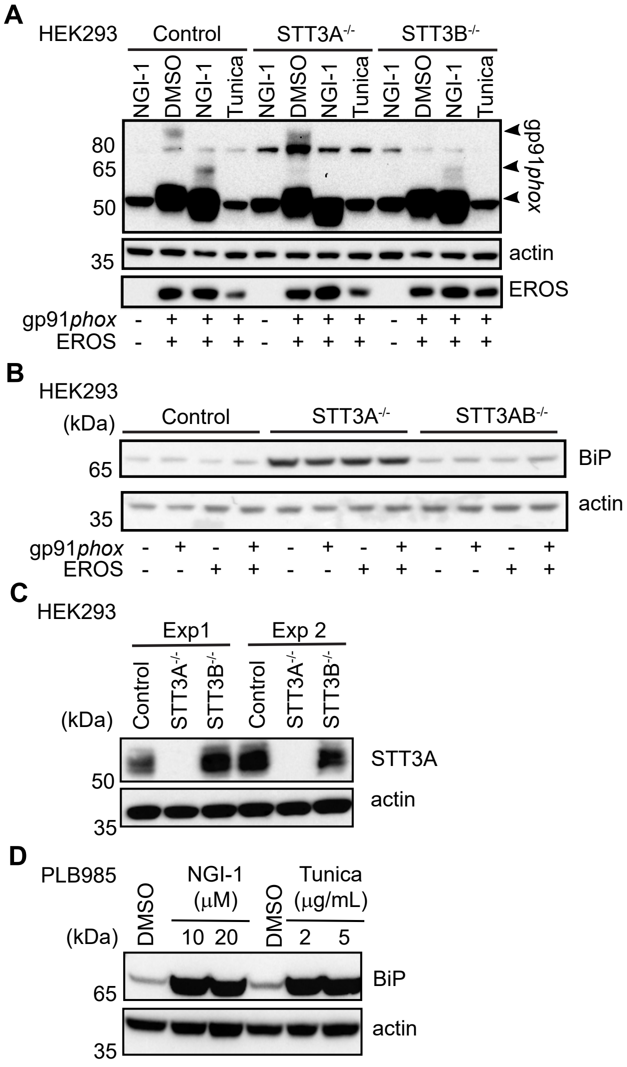

### Figure 5- figure supplement 1

**Figure 5 - Figure supplement 1**

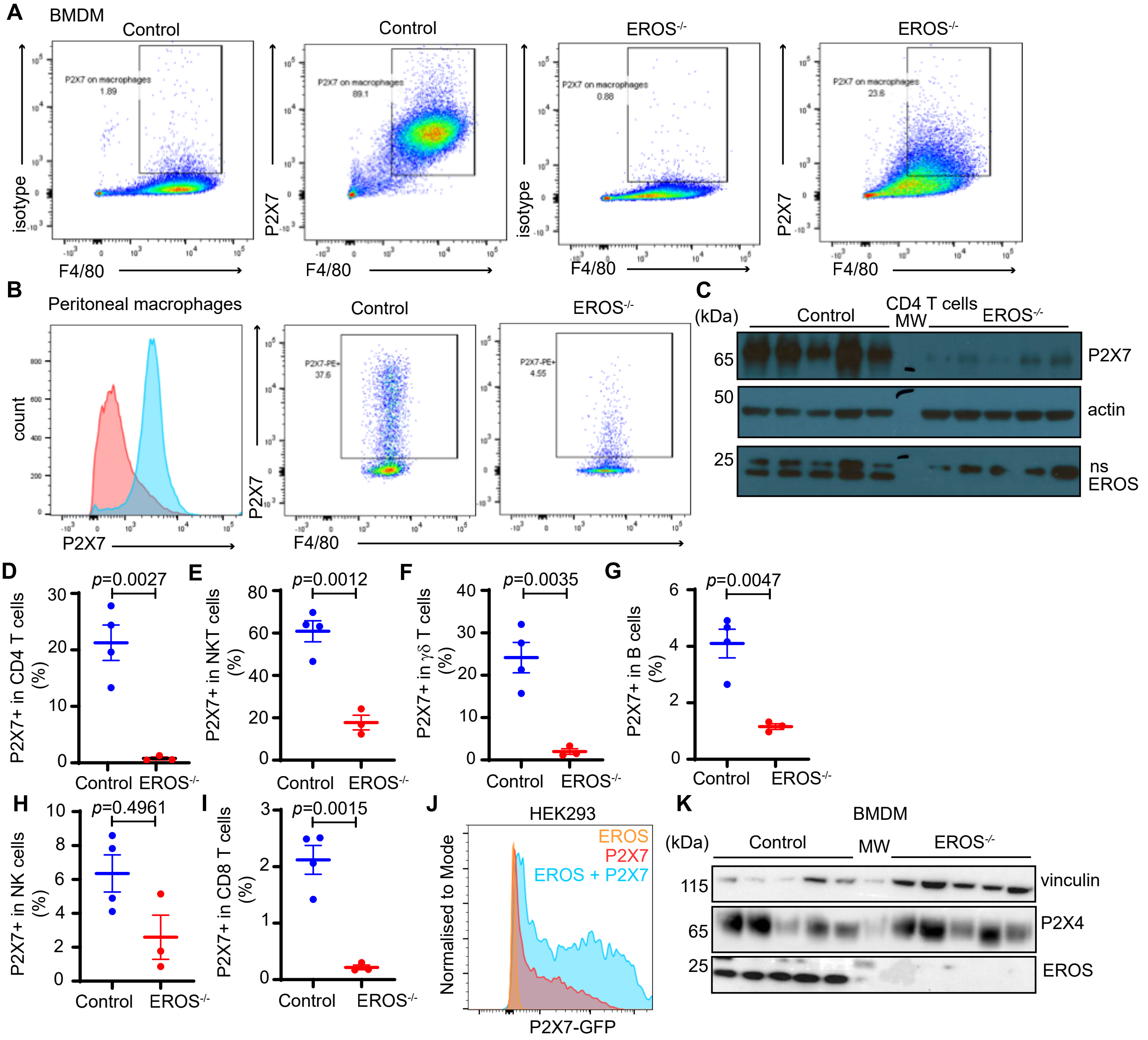

### Figure 6- figure supplement 1

**Figure 6 - Figure supplement 1**

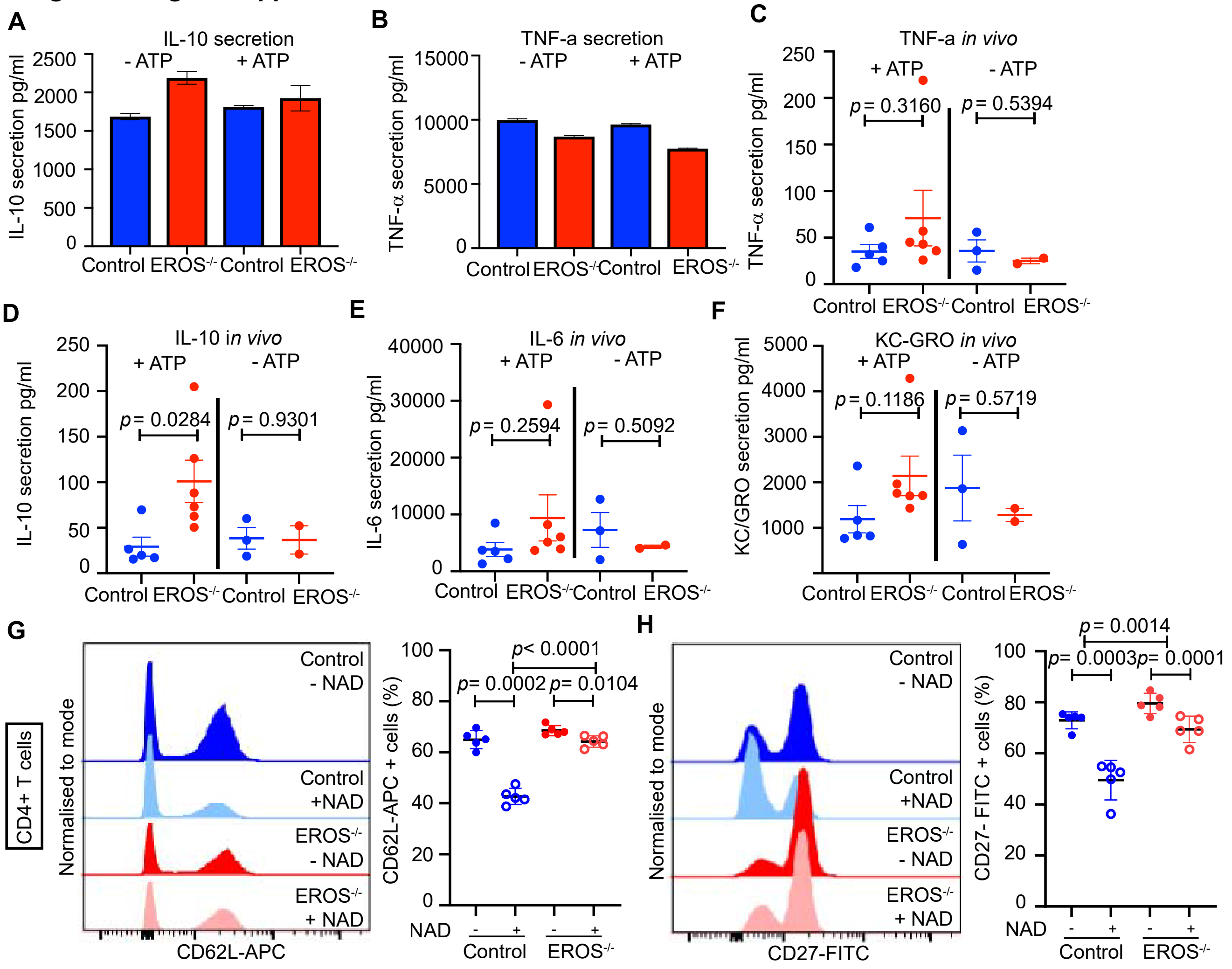
